# A CandiChrome toolkit for multicolor labeling of *Candida* cells

**DOI:** 10.64898/2026.05.11.723596

**Authors:** Cai-Ling Ke, Jingcheng Xu, Corey Frazer, Richard J. Bennett

## Abstract

Here, we develop CandiChrome, a multiplex labeling toolkit for *Candida albicans*, through combined *in vitro* and *in vivo* characterization of fluorescent proteins in a standard strain background. To this end, we screened 13 candidate fluorophores across the visible spectrum and assessed their practical performance based on brightness, stability, and usability. This analysis identified a seven-fluorophore set that achieved the most effective balance of signal strength, robustness, and compatibility. We used this optimized panel to build a modular multicolor platform that enables strain labeling, mixed-population imaging, and competition assays in *C. albicans*. This platform could resolve up to 21 distinct populations by flow cytometry and microscopy. Importantly, CandiChrome supported the resolution of differentially labeled populations both *in vitro* and in the murine host, supporting the simultaneous tracking of multiple strains in complex settings. Together, these results establish CandiChrome as a flexible platform for multiplex fungal imaging in a pathogenic species where multicolor tools remain underdeveloped.

## Introduction

Fluorescent protein-based labeling has become an essential approach for visualizing protein localization and for monitoring population dynamics in living systems [1]. Fluorescent protein toolkits can include a broad spectral range of fluorophores, enabling simultaneous observation of multiple biological features in the same sample [1–12]. In parallel, combinatorial multicolor strategies such as Brainbow demonstrated that stochastic expression of multiple fluorophores can increase the power of imaging-based discrimination within complex cell populations [13]. Nevertheless, limitations exist because of spectral overlap, unequal signal intensity, and the need for system-specific implementation. For example, a *Drosophila* adaptation of Brainbow identified five compatible colors that were suitable for combinatorial neural imaging [13]. More recently, multiplex fluorescent labeling has been enabled by computational unmixing, increasing the number of resolvable labels beyond what can typically be achieved by standard microscopy [14]. Notably, Tetbow enabled seven-color labeling of layer 2/3 neurons in the primary somatosensory cortex [14]. Together, these studies highlight the potential of multiplex labeling for dissecting heterogeneous biological systems.

Despite this progress, the implementation of multi-label imaging strategies for new targets has several potential hurdles. First, fluorophores with attractive properties do not always perform equivalently in different cellular environments because effective brightness depends not only on intrinsic parameters but also on folding efficiency, maturation, degradation, pH sensitivity, and photostability [15]. Indeed, many fluorescent proteins expressed in the budding yeast *Saccharomyces cerevisiae* behave differently from that expected based on their behavior in other systems, underscoring the need for species-specific evaluation [15]. Second, spectral overlap, photochromism, and unequal expression or maturation rates can complicate multi-color imaging, especially when several fluorophores are combined in the same biological system. As evident in neuronal labeling systems, increasing the number of fluorophores beyond three introduces challenges in visual interpretation and often requires dedicated computational pipelines for signal separation and color assignment. These issues are also relevant to fungal species where fluorescent labeling platforms are often less developed than those in higher eukaryotes. In *S. cerevisiae*, substantial efforts have been made to generate fluorescent protein marker sets, providing a resource for live-cell imaging [16]. In *Candida albicans*, fluorescent reporters have been utilized for gene expression analysis, protein localization, and cell state identification, including early GFP-based reporters [17, 18], fluorescent protein tagging tools [19, 20], codon-optimized fusion cassettes [20], and photostable GFP variants [21]. More recently, *Candida*-optimized next-generation fluorophores such as mNeonGreen and mScarlet were shown to outperform GFP- and mCherry-based reporters in specific applications [22]. However, there remains a limited set of fluorophores for use in *C. albicans* despite its importance as a prevalent human commensal and opportunistic pathogen.

Here, we developed a multicolor platform for *C. albicans* that is ready to implement for competition assays, mixed-population analyses, and other applications requiring simultaneous tracking of multiple strains or proteins. These efforts led us to establish CandiChrome as a modular multicolor labeling platform for *C. albicans*.

## Results

### CandiChrome for multicolor labeling of *Candida* cells

To enable multicolor labeling of *C. albicans* cells, we selected fluorescent proteins with distinct excitation and emission spectra, and included candidates with reported advantages in brightness, photostability, or spectral separability (Fig. 1A). For example, mTagBFP2 was selected as a strong blue fluorescent protein [14, 23], whereas mTurquoise2 and mAmetrine1.1 were included because prior spectral analyses suggested that they might be distinguishable from mTagBFP2 and from other fluorophores in multiplexed applications [14]. Among green fluorescent proteins, mStayGold was included for comparison with the commonly used mNeonGreen and mEGFP, given the report of higher fluorescence intensity and improved photostability [22, 24]. For red fluorophores, dTomato and mScarlet-I were included as widely used fluorophores, whereas mScarlet3-S2 was selected as an alternative to mScarlet-I because of its reported greater photostability [7, 8, 22, 25]. mKate2 was also included following prior work showing brightness and photostability comparable to dTomato in budding yeast [15]. To extend the panel beyond the visible spectrum, we selected miRFP670nano3, BDFP1.6, and smURFP as far-red and near-infrared candidates on the basis of their brightness in mammalian systems and their spectral separation from visible-range fluorophores [11, 26, 27]. Target fluorophores were optimized and synthesized for expression in *C. albicans* and compared to those previously described [22]. In total, 13 fluorophores were evaluated by expression as a C-terminal fusion with *C. albicans* Eno1 at the native *ENO1* locus (Fig. 1B). Labeled strains grew at similar rates to the unlabeled parental strain (SC5314) in four different media, indicating that fluorophore expression did not affect overall growth (Fig. 1C).

**Figure 1.**
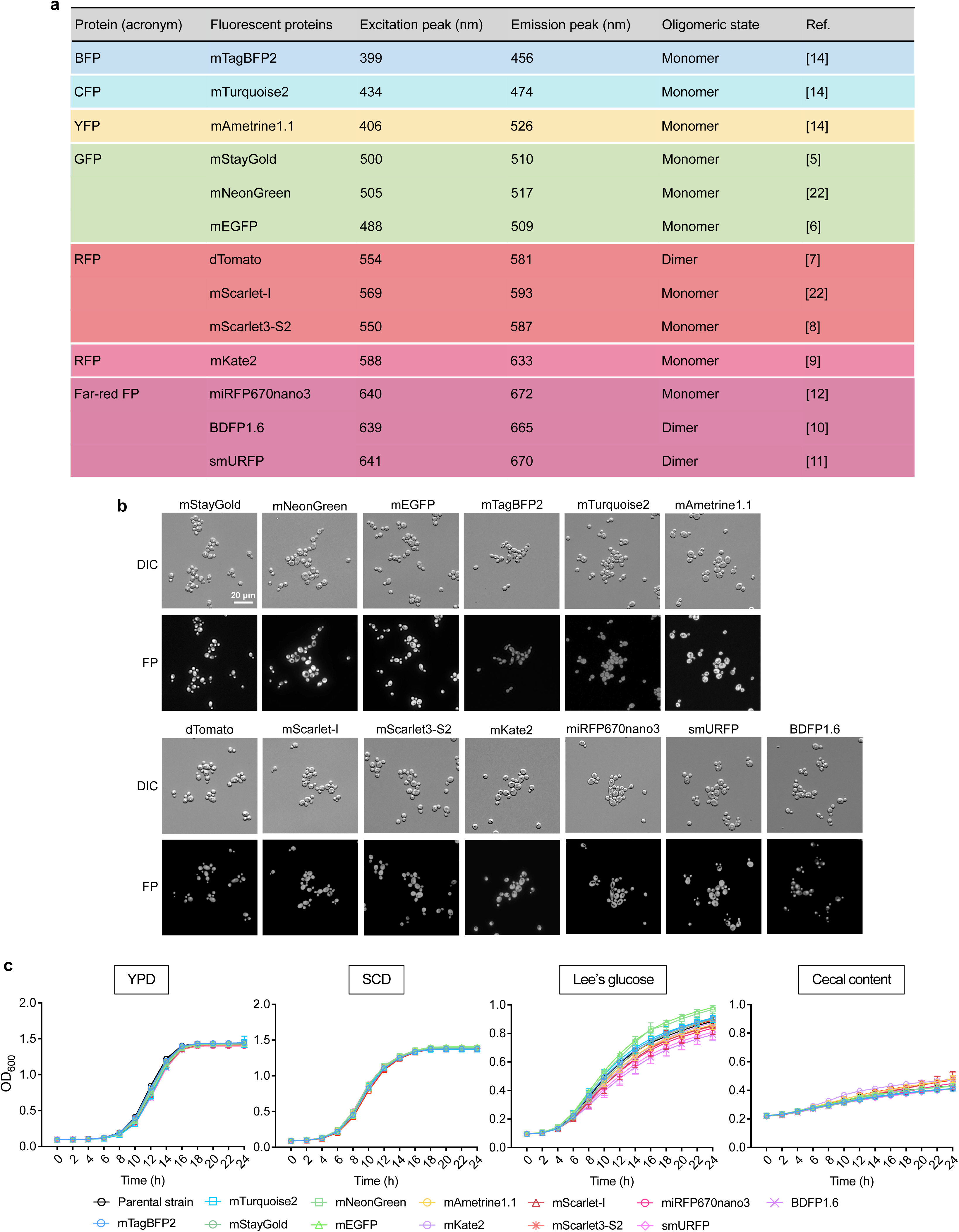
Candidate fluorescent proteins for the CandiChrome panel. **a,** Thirteen candidate fluorescent proteins spanning the blue-to-far-red spectrum were tested in *C. albicans*. **b,** Representative DIC and fluorescence images of *C. albicans* strains expressing the indicated Eno1–fluorophore fusions acquired using an Evident APEXVIEW APX100 microscope with DAPI, GFP, Cy3, and Cy5 imaging channels. **c,** Growth of strains expressing the indicated strains expressing different Eno1–FPs when cultured at 30°C in YPD, SCD, Lee’s glucose, or cecal content media. Optical density at 600 nm (OD_600_) was measured over time. Data are shown as mean ± s.d. of biological replicates.

### Comparison of fluorescence intensity and photostability of CandiChrome fluorescent proteins

We compared the fluorescence intensity and stability of each protein by both microscopy and flow cytometry to identify those suitable for multicolor labeling in *C. albicans*. For microscopy, we used an APEXVIEW APX100 microscope (Evident Scientific) equipped with DAPI, GFP, Cy3, and Cy5 filter cubes. We grouped the selected fluorescent proteins according to their excitation spectra, which broadly corresponded to their fluorescence color. Among green fluorescent proteins, mEGFP exhibited the highest fluorescence intensity followed by mNeonGreen, whereas mStayGold showed low fluorescence intensity (Fig. 2A). Analysis showed that all three green fluorescent proteins displayed similar photostability (Fig. 2B). Consistent with these results, flow cytometry showed that mEGFP had the highest intensity among the green fluorescent proteins (Fig. 2C).

**Figure 2.**
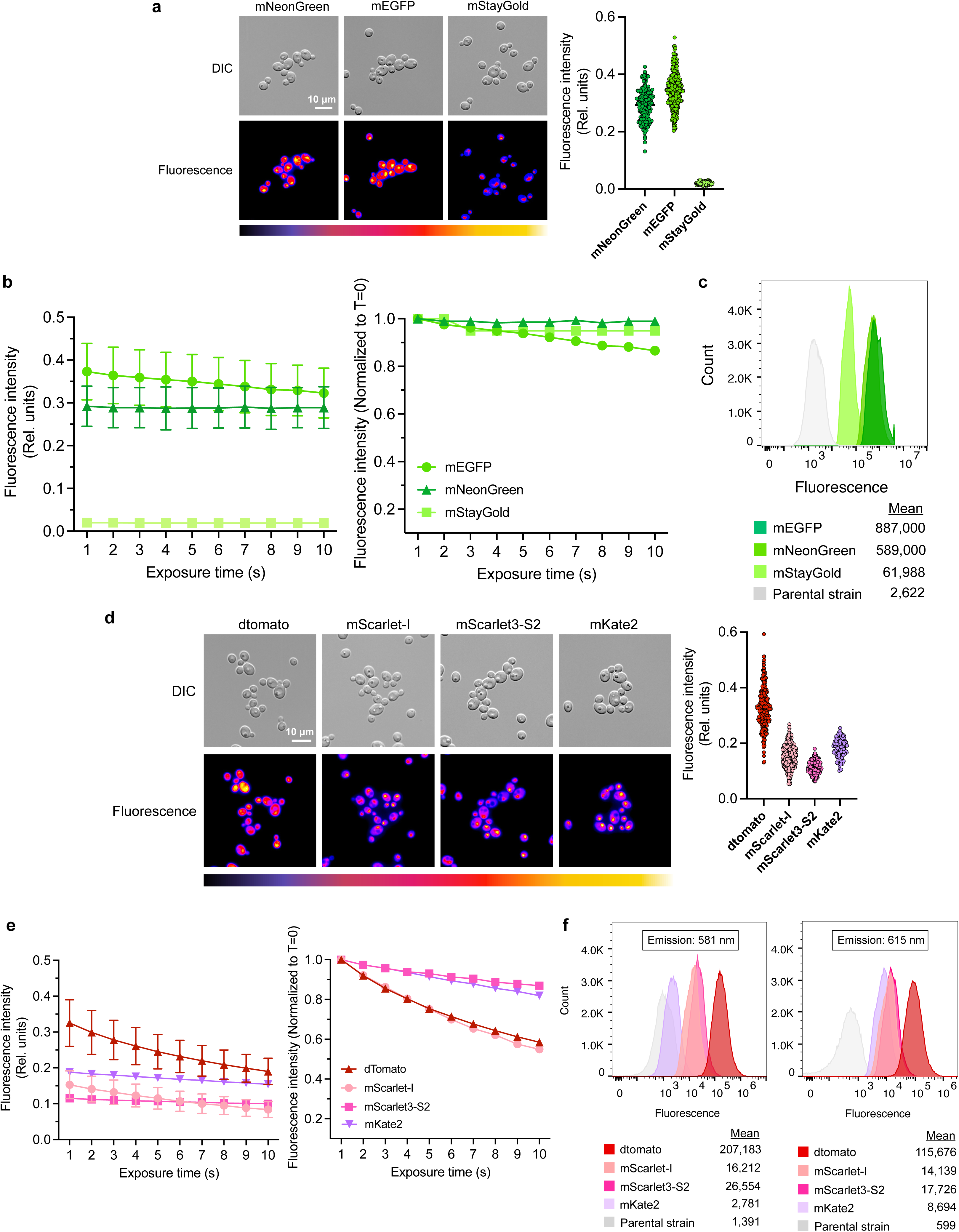
Brightness and photostability of fluorescent proteins in *C. albicans*. Cells were grown overnight in SCD at 30 °C before analysis. Fluorescence microscopy images were acquired on an Evident APEXVIEW APX100 microscope using the GFP or Cy3 imaging channels, as indicated for each panel. Flow cytometry data were acquired on a Cytek Aurora flow cytometer equipped with UV, violet, blue, and red lasers. **a,** Representative DIC and fluorescence images of strains expressing Eno1-mNeonGreen, Eno1-mEGFP, or Eno1-mStayGold are shown (left). Single-cell average pixel intensity was quantified using CellProfiler from at least 200 cells per strain (right). Error bars represent s.d.; Rel. units, relative units. **b,** Photostability was examined by repeated 1 s exposures at 30% light intensity on an Evident APX100 microscope using the GFP imaging channel. Fluorescence intensity values (left) and values normalized to T = 0 (right) are shown. Average pixel intensity was measured from at least 200 cells per strain in a representative field of view. Error bars represent s.d.; Rel. units, relative units. **c,** Histograms show fluorescence intensity distributions of 100,000 events acquired on an Aurora flow cytometer, with fluorescence detected in the 508 nm emission channel. Mean fluorescence intensity values are indicated below the histogram. **d,** Representative DIC and fluorescence images of strains expressing Eno1-dTomato, Eno1-mScarlet-I, Eno1-mScarlet3-S2 or Eno1-mKate2 are shown (left). Fluorescence images were acquired on an Evident APX100 microscope using the Cy3 channel. Single-cell average pixel intensity was quantified using CellProfiler from at least 200 cells per strain (right). Error bars represent s.d.; Rel. units, relative units. **e,** Cells were photobleached on an Evident APX100 microscope in the Cy3 channel by repeated 1 s exposures at 30% light intensity. Fluorescence intensity values (left) and values normalized to T = 0 (right) are shown. Average pixel intensity was measured from at least 200 cells per strain in a representative field of view. Error bars represent s.d.; Rel. units, relative units. **f,** Histograms show fluorescence intensity distributions of 100,000 events acquired on an Aurora flow cytometer and detected in the 581 nm and 615 nm emission channels. Mean fluorescence intensity values are indicated below the histograms.

Among red fluorescent proteins, dTomato was the brightest by microscopy (Fig. 2D). and remained the brightest after 10 seconds of photobleaching, even though dTomato and mScarlet-I showed greater relative signal loss than mScarlet3-S2 and mKate2 over the bleaching period (Fig. 2E). We also evaluated red fluorescence by flow cytometry using two detection channels; a 581-nm emission setting is more suitable for most red fluorescent proteins whereas mKate2 is better detected at 615 nm. Consistent with the microscopy results, dTomato exhibited the highest fluorescence intensity in both flow cytometry channels (Fig. 2F).

Among far-red fluorescent proteins, BDFP1.6 and miRFP670nano3 showed comparable intensity by microscopy whereas smURFP was much lower intensity (Fig. 3A). BDFP1.6, miRFP670nano3, and smURFP were all highly photostable and resisted photobleaching (Fig. 3B). Consistent with the microscopy results, flow cytometry showed comparable fluorescence intensity for BDFP1.6 and miRFP670nano3 with smURFP being much weaker (Fig. 3C).

**Figure 3.**
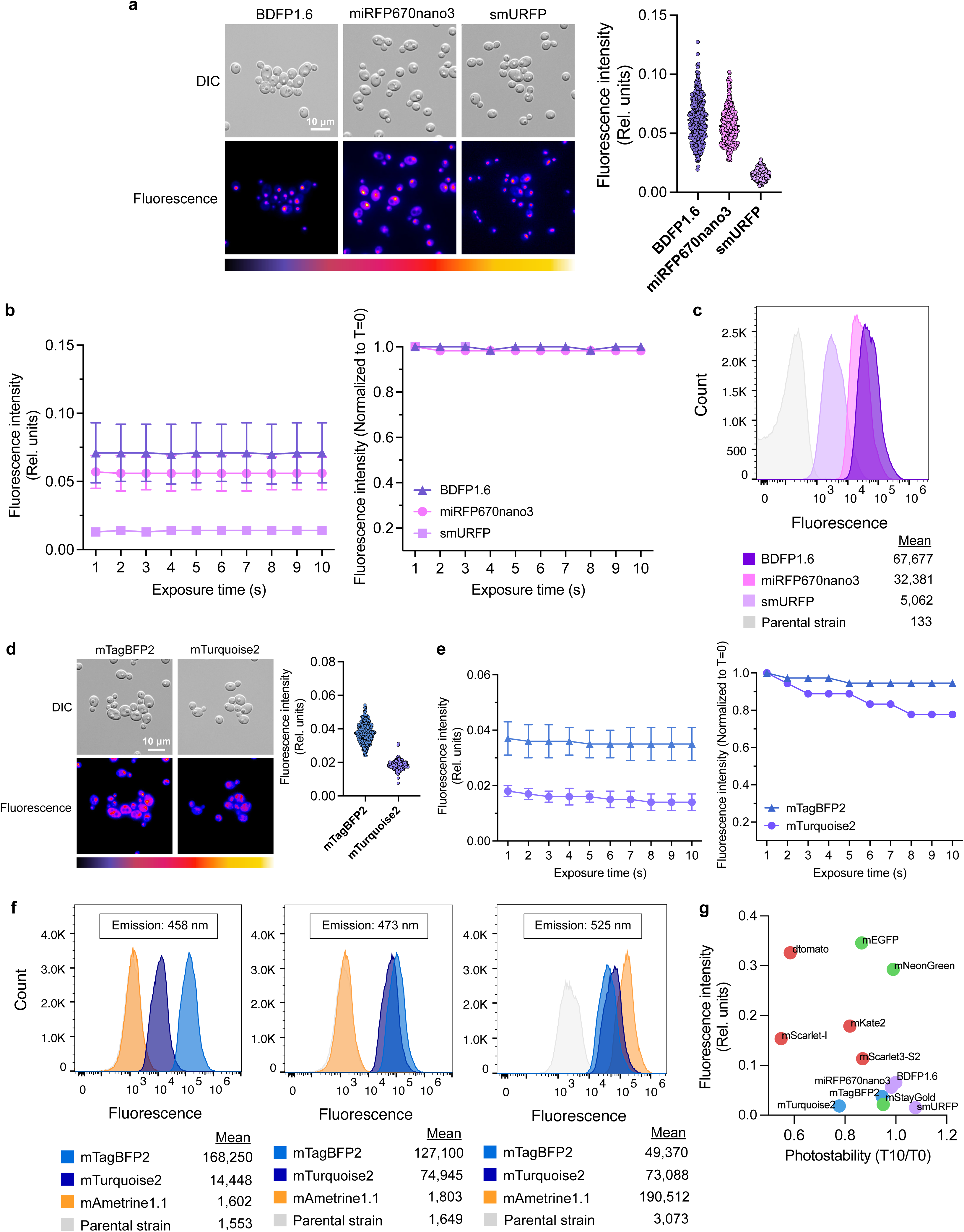
Evaluation of additional fluorescent proteins in *C. albicans*. Cells were grown overnight in SCD at 30 °C before analysis. Fluorescence microscopy images were acquired on an Evident APEXVIEW APX100 microscope using the Cy5 or DAPI imaging channels, as indicated. Flow cytometry data were acquired on a Cytek Aurora flow cytometer equipped with UV, violet, blue, and red lasers. **a,** Representative DIC and fluorescence images of strains expressing Eno1-BDFP1.6, Eno1-miRFP670nano3 and Eno1-smURFP are shown (left). Fluorescence images were acquired using the Cy5 channel. Single-cell average pixel intensity was quantified using CellProfiler from at least 200 cells per strain (right). Error bars, s.d.; Rel. units, relative units. **b,** Cells were progressively photobleached in the Cy5 channel by repeated 1 s exposures at 30% light intensity. Fluorescence intensity values (left) and values normalized to T = 0 (right) are shown. Average pixel intensity was measured from at least 200 cells per strain in a representative field of view. Error bars, s.d.; Rel. units, relative units. **c,** Histograms show fluorescence intensity distributions of 100,000 events acquired on an Aurora flow cytometer and detected in the 661 nm emission channel. Mean fluorescence intensity values are shown below the histogram. **d,** Representative DIC and fluorescence images of strains expressing Eno1-mTagBFP2 or Eno1-mTurquoise2 are shown (left). Fluorescence images were acquired using the DAPI imaging channel. Single-cell average pixel intensity was quantified using CellProfiler from at least 200 cells per strain (right). Error bars, s.d.; Rel. units, relative units. **e,** Cells were progressively photobleached in the DAPI channel by repeated 1 s exposures at 30% light intensity. Fluorescence intensity values (left) and values normalized to T = 0 (right) are shown. Average pixel intensity was measured from at least 200 cells per strain in a representative field of view. Error bars, s.d.; Rel. units, relative units. **f,** Histograms show fluorescence intensity distributions of 100,000 events acquired on an Aurora flow cytometer in the 458 nm, 473 nm and 525 nm emission channels. Mean fluorescence intensity values are shown below the histograms. **g,** Fluorescence intensity and photostability values for the indicated fluorescent proteins are plotted to summarize relative performance across the candidate panel. Rel. units, relative units.

In the absence of a suitable filter cube for mAmetrine1.1, only mTagBFP2 and mTurquoise2 were evaluated by microscopy using the DAPI channel (Fig. 3D). Under these conditions, mTagBFP2 exhibited higher fluorescence intensity than mTurquoise2, likely reflecting its greater compatibility with the DAPI filter. Analysis further showed that mTagBFP2 was more photostable than mTurquoise2 (Fig. 3E). We compared all three proteins by flow cytometry using three detection channels corresponding to 458 nm, 473 nm, and 525 nm, which are optimal for mTagBFP2, mTurquoise2, and mAmetrine1.1, respectively. In the 458-nm and 473-nm channels, mTagBFP2 showed the highest overall intensity (although the mTurquiose2 signal was similar to the mTagBFP2 signal in the 473-nm channel) while mAmetrine1.1 exhibited the highest intensity in the 525-nm channel, consistent with its suitability to that channel (Fig. 3F).

To identify the fluorescent proteins most suitable for simultaneous use in a multicolor labeling platform, we compared all candidates based on both fluorescence intensity and photostability (Fig. 3G). Among the 13 proteins tested, mEGFP, mNeonGreen, and dTomato showed the best overall performance, with mKate2, mScarlet-I and mScarlet3-S2 in the next strongest group, and with the remaining proteins (mTurquoise2, mTagBFP2, miRFP670nano3, BDFP1.6, mStayGold and smURFP) showing the weakest fluorescence intensity.

### Identification of a CandiChrome panel for multicolor labeling of *C. albicans*

Based on the results described above, a fluorescent panel for combinatorial seven-color labeling in *C. albicans* was developed and termed CandiChrome. This panel consists of mTagBFP2, mTurquoise2, mAmetrine1.1, mEGFP/mNeonGreen, dTomato/mScarlet-I, mKate2, and miRFP670nano3 (Fig. 4A). To avoid potential confounding effects of the expression of fluorophores fused to Eno1, each fluorophore was evaluated in its native form when expressed under the *pENO1* promoter at neutral genetic locus, that of *C. albicans NEUT5L* [28, 29].

**Figure 4.**
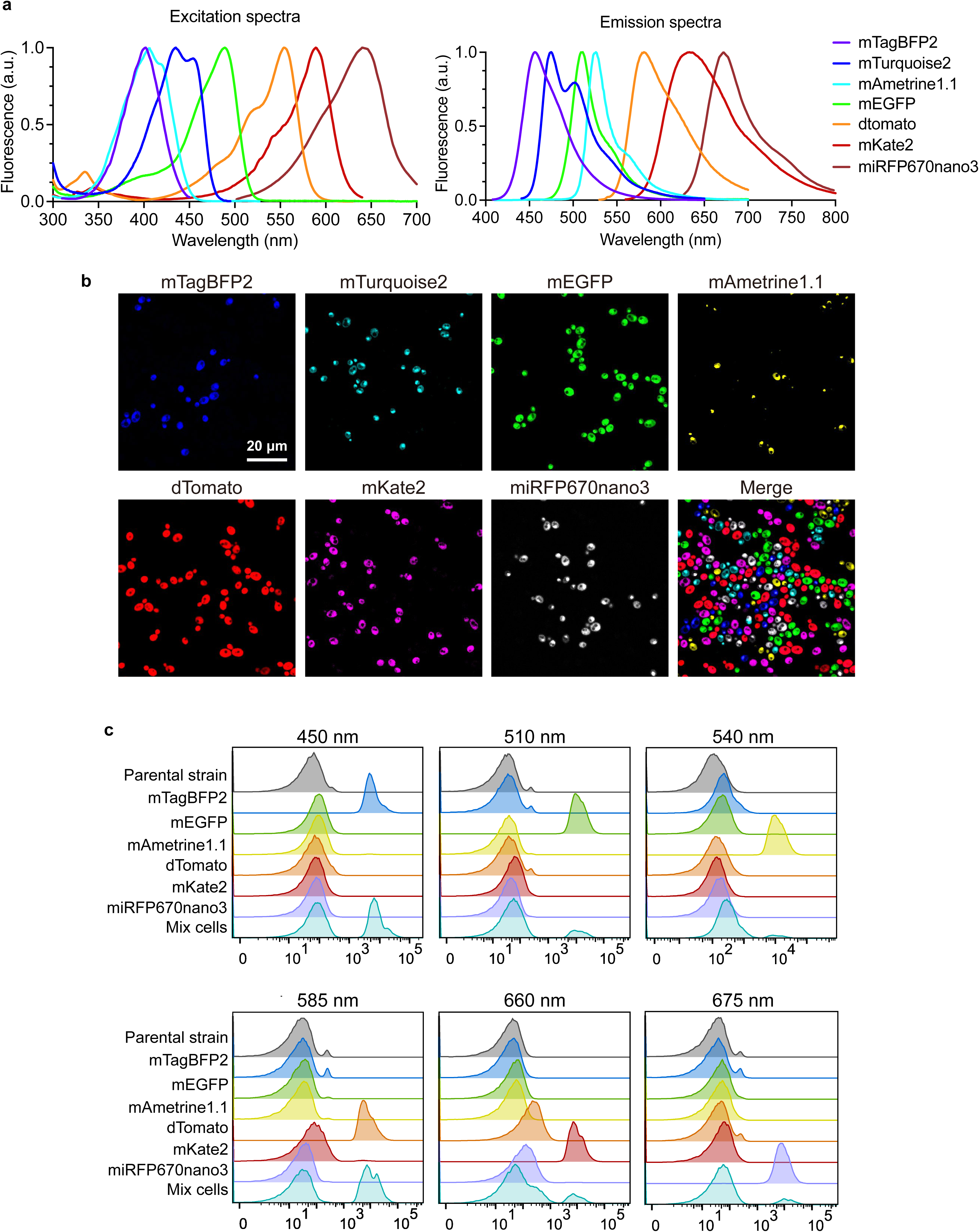
Multicolor imaging and flow cytometric resolution with CandiChrome. **a,** Excitation and emission spectra of the seven fluorescent proteins selected for CandiChrome: mTagBFP2, mTurquoise2, mAmetrine1.1, mEGFP, dTomato, mKate2 and miRFP670nano3. **b,** Representative fluorescence images of *C. albicans* strains expressing the indicated fluorophores under the *pENO1* promoter at the *NEUT5L* locus. Cells were grown overnight in SCD at 30 °C and imaged on an FV5000 confocal microscope with 405, 445, 488, 561, 594, and 640 nm lasers. **c,** Flow cytometry analysis of individual and mixed strains expressing the six fluorophores, acquired on a FACSymphony flow cytometer with violet, blue, yellow-green, and red laser lines.

We validated the CandiChrome panel by microscopy and flow cytometry. For microscopy-based evaluation, labeled strains were imaged using a FLUOVIEW FV5000 confocal microscope (Evident) equipped with excitation lasers matched to the spectral properties of the selected proteins. Specifically, 405 nm was used for mTagBFP2 and mAmetrine1.1, 445 nm for mTurquoise2, 488 or 514 nm for mEGFP/mNeonGreen, 561 nm for dTomato/mScarlet-I, 594 nm for mKate2, and 640 nm for miRFP670nano3. As shown in Fig. 4B, each of the fluorescent signals were distinguishable, demonstrating that CandiChrome enables simultaneous seven-color imaging in *C. albicans*. We also tested the same strains on a FLUOVIEW FV3000 confocal microscope (Evident) equipped with four excitation lasers (405, 488, 561, and 640 nm). Here, separation of the fluorescent signals required spectral demixing which nevertheless supported discrimination of the seven labeled populations (Supplementary Fig. S1).

This multicolor panel was also analyzed using a BD FACSymphony A5 SE flow cytometer equipped with five lasers (349, 404, 488, 561, and 637 nm). The emission spectra of mTagBFP2 and mTurquoise2 were too close to allow reliable separation and so these fluorophores could not be distinguished from each other. Nevertheless, the remaining six fluorescent signals still allowed six-color labeling of *C. albicans* populations by flow cytometry. Both singly labeled samples and a mixed sample containing all labeled strains were analyzed. Following spectral unmixing using the instrument’s built-in software, the distinguishable fluorophores were well separated, and the mixed sample showed clear separation of positive and negative populations for each fluorophore (Fig. 4C). Together, these results demonstrate that CandiChrome provides a suitable platform for six-color flow cytometric analysis in *C. albicans*.

### Double-color labeling expands the CandiChrome platform

After establishing that six CandiChrome fluorophores could be distinguished by both flow cytometry and microscopy, we next generated double-labeled strains to increase the number of populations that can be distinguished. Including the original single-labeled strains, combinations of the six fluorophores yielded a total of 21 *C. albicans* populations for analysis. We mixed the 21 samples at equal ratios and analyzed them using a Cytek Aurora cytometer (Cytek Biosciences). The resulting data were subjected to automated deconvolution and population quantification followed by UMAP generation in R. As shown in Fig. 5A, all 21 populations were well separated, and the observed proportions closely matched the input ratios. We also analyzed a mixture containing only 16 samples using the same workflow. In this case, UMAP analysis again showed clear separation of each population, whereas populations not included in the mixture were not represented in the UMAP distribution (Supplementary Fig. S2). The 21-sample mixture was also evaluated on an Evident FV5000 confocal microscope. Examination of the individual channel images allowed representative cells corresponding to each fluorescent combination to be identified, supporting the visual distinguishability of the selected fluorophore combinations (Fig. 5B).

**Figure 5.**
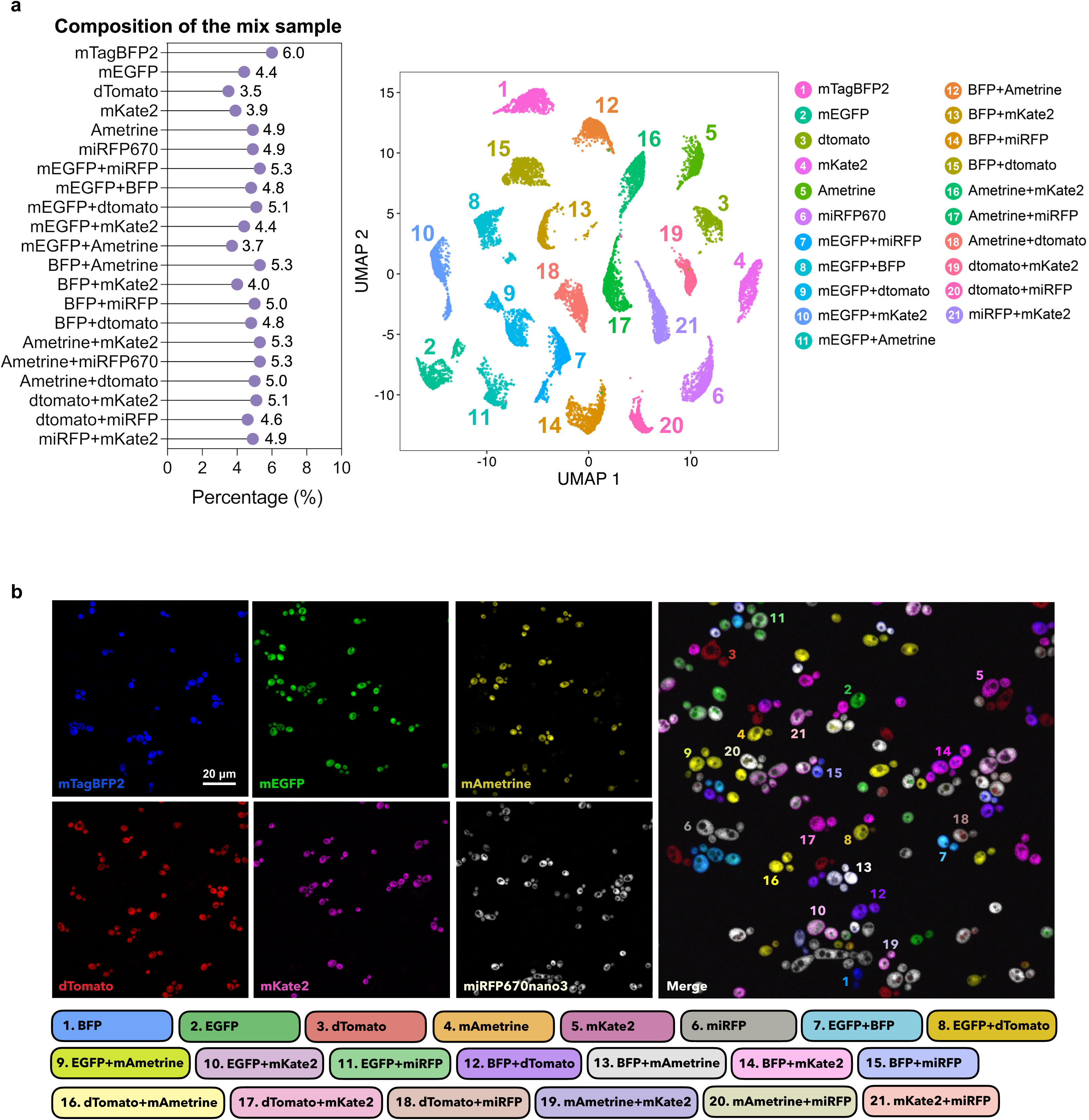
Resolution of dual-color CandiChrome-labeled populations. Cells were grown overnight in SCD at 30 °C before analysis. All strains expressed FPs under the *pENO1* promoter at the *NEUT5L* locus. **a,** Composition of the mixed sample is shown (left), and UMAP projection of flow cytometry data is shown (right). Data were acquired on a Cytek Aurora cytometer and analyzed in R. Distinct clusters corresponding to the indicated single- and dual-labeled strains were identified. **b,** Representative fluorescence images of the indicated single-color channels and the merged image were acquired on a FV5000 microscope. Numbers in the merged image correspond to the labeled populations listed below.

### Colony-level imaging resolves a subset of the CandiChrome panel

To assess whether a subset of the CandiChrome panel could be detected using a colony-level imaging workflow, plates were imaged using a ChemiDoc MP Imaging System (Bio-Rad). Under these conditions, three acquisition channels detected a subset of the fluorophores at the colony level: Alexa488 detected mEGFP and mNeonGreen, Cy3 detected dTomato, mScarlet-I, and mKate2, and Cy5 detected miRFP670nano3 (Fig. 6A–C). In contrast, mTagBFP2 (Fig. 6A) and mAmetrine (Fig. 6B) did not produce a detectable signal and thus these colonies were similar to the unlabeled parental control (Fig. 6C). These results show that a subset of the CandiChrome panel can be resolved at the colony level using a standard gel imaging system.

**Figure 6.**
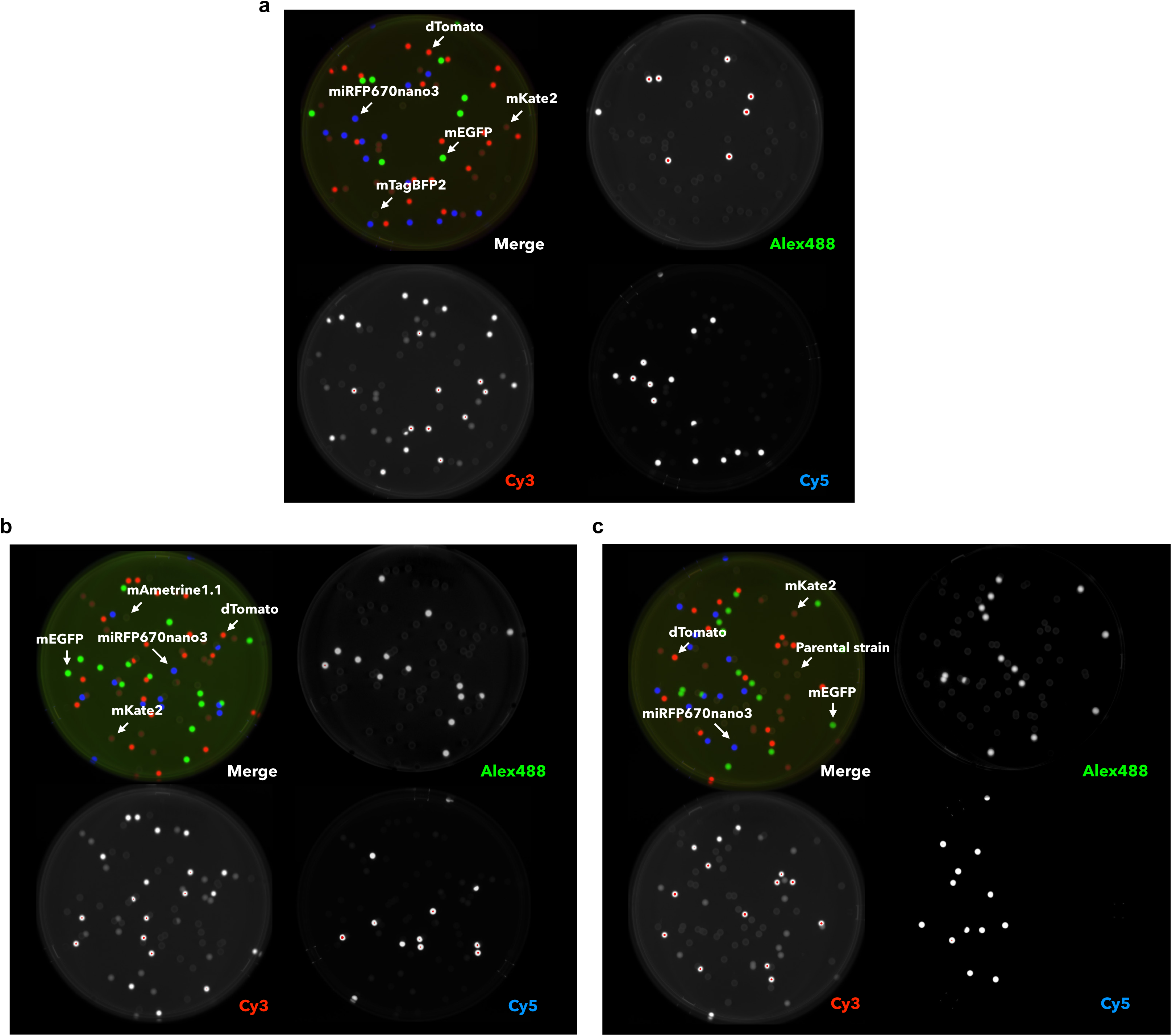
Direct colony imaging with fluorescently labeled cells. **a–c,** Representative images of *C. albicans* strains expressing the indicated CandiChrome fluorophores at the *NEUT5L* locus. Images of colonies were acquired using a ChemiDoc MP Imaging System (Bio-Rad) in the Alexa488, Cy3 and Cy5 channels. **a,** Mixture of strains expressing mTagBFP2, mEGFP, dTomato, mKate2 and miRFP670nano3. **b,** Mixture of strains expressing mAmetrine1.1, mEGFP, dTomato, mKate2 and miRFP670nano3. **c,** Mixture of strains expressing mEGFP, dTomato, mKate2, miRFP670nano3 and an unlabeled control.

### CandiChrome enables multicolor competition assays in the murine host

To assess the applicability of CandiChrome for *in vivo* competition assays, we co-inoculated six fluorescently labeled *C. albicans* strains into five antibiotic-treated C57BL/6J mice (Comp1-Comp5). Mice received drinking water supplemented with penicillin (1.5 mg/mL), streptomycin (2 mg/mL), and 2.5% glucose throughout the experiment (Fig. 7A). To establish controls for spectral demixing, parental and singly labeled strains were also inoculated individually into mice. All groups established robust gut colonization, with fungal levels in fecal pellets ranging from approximately 10^7^ to 10^8^ cells/g (Fig. 7A). Competition samples recovered from fecal pellets were plated on YPD and incubated at 30°C for 3 days. Colonies were then collected from the agar and analyzed on a BD FACSymphony flow cytometer. The relative abundance of each population remained relatively stable, at levels close to the initial input ratio (Fig. 7B), although significant differences were detected between the mEGFP- and mAmetrine-labeled strains and between the miRFP670nano3- and mAmetrine-labeled strains at days 7 and 10 after inoculation. These differences did not persist and were absent from later time points and from *C. albicans* cells recovered from GI organs (Fig. 7B and Supplementary Fig. S3). Importantly, different fluorescent signals remained distinguishable in samples recovered from the *in vivo* competition experiments. A representative flow cytometry plot of an *in vivo* sample is shown in Supplementary Fig. S4. These results demonstrate that CandiChrome can be effectively applied to *in vivo* competition assays in *C. albicans*.

**Figure 7.**
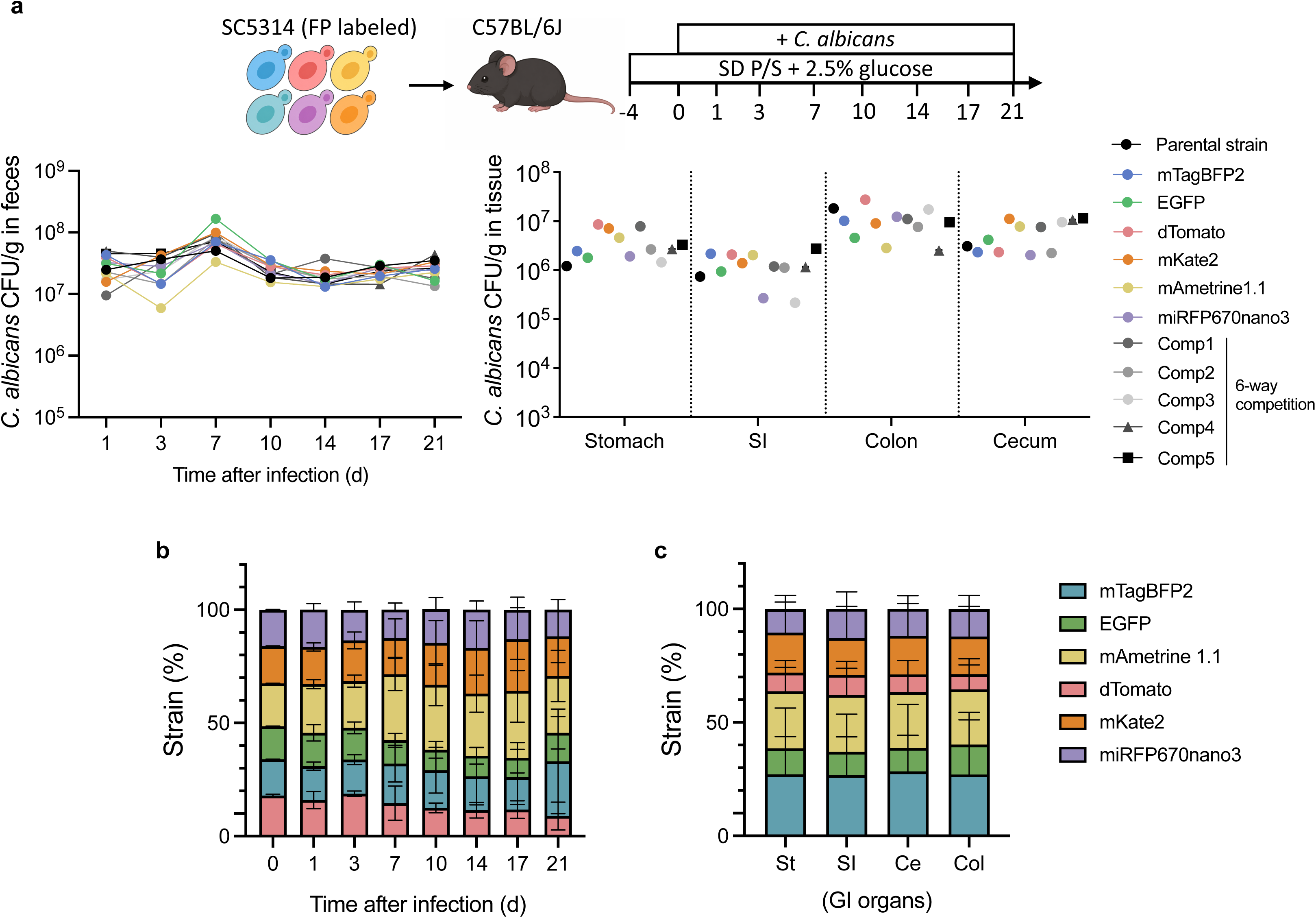
*In vivo* performance of CandiChrome-labeled populations in a mouse gastrointestinal colonization model. **a,** Schematic of the experimental design (top). C57BL/6J mice were gavaged with a mixed population of fluorescently labeled *C. albicans* strains and received antibiotics with 2.5% glucose throughout the experiment. Fecal fungal burdens were measured over time as *C. albicans* CFU g^−1^ feces (bottom left), and tissue fungal burden was measured at endpoint in the indicated gastrointestinal sites (bottom right). Symbols indicate individual mice. **b,** Stacked bar plots show the proportion of each fluorescent strain recovered at the indicated time points. Data are shown as mean ± s.d. **c,** Stacked bar plots show the proportion of each strain recovered from stomach (St), small intestine (SI), cecum (Ce) and colon (Col). Data are shown as mean ± s.d.

### CandiChrome supports protein localization and expression studies in *C. albicans*

Beyond strain tracking, CandiChrome can be applied to protein localization and expression analysis in *C. albicans*. To illustrate this versatility, we used mTagBFP2 to label the nuclear transcription factor Efg1, dTomato to label the mitochondrial factor Cox4, miRFP670nano3 to label the vacuolar protein Vph1, and mNeonGreen to label the plasma membrane-localized protein Ras1. As shown in Fig. 8, each fusion protein exhibited the expected subcellular localization pattern, demonstrating that CandiChrome can be used to simultaneously visualize multiple cellular proteins/compartments in *C. albicans*. Together, these results establish CandiChrome as a versatile multicolor labeling platform for protein localization studies and *in vivo* competition assays.

**Figure 8.**
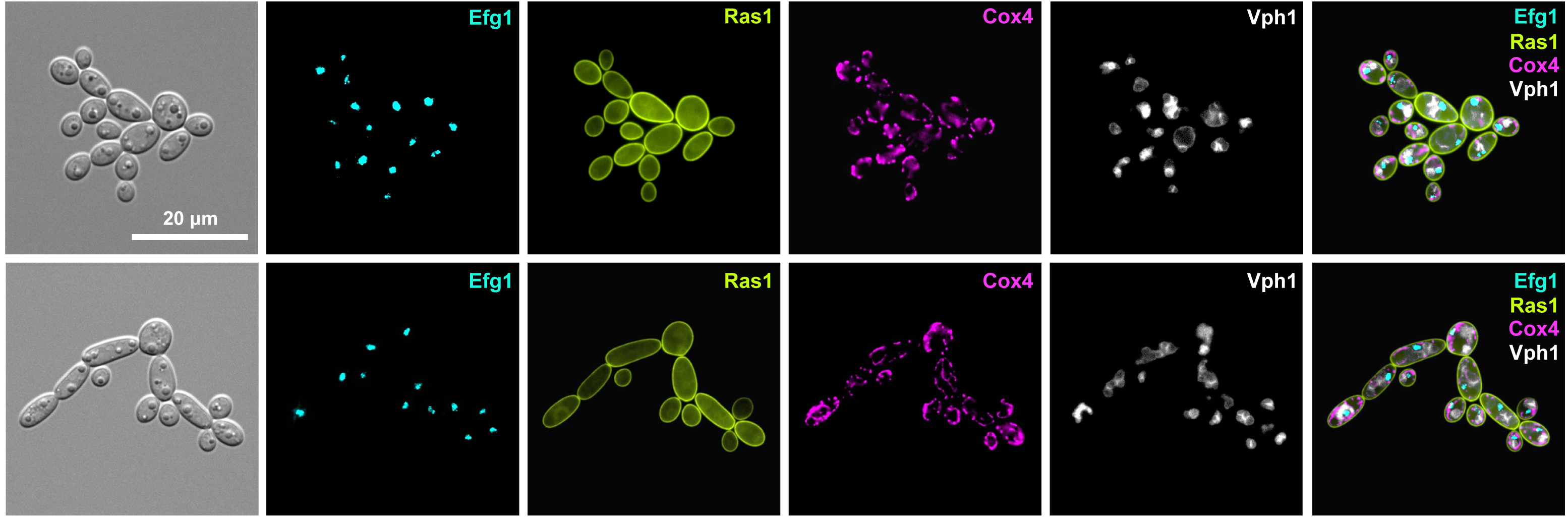
Multicolor protein localization with CandiChrome in *C. albicans*. Representative images of cells expressing Efg1-mTagBFP2, Ras1-mNeonGreen, Cox4-dTomato and Vph1-miRFP670nano3. Cells were grown overnight in RPMI 1640 medium at 30 °C and imaged on an Evident APX100 microscope using the DAPI, GFP, Cy3, and Cy5 imaging channels.

## Discussion

Multiplex fluorescent labeling approaches have greatly expanded the ability to resolve complex cellular populations in many experimental systems, yet comparable tools in *C. albicans* have remained limited. Here, we establish CandiChrome, a multicolor fluorescent labeling platform for *C. albicans* through systematic evaluation of fluorescent proteins in a standard strain background.

By screening 13 candidate fluorophores and selecting a seven-fluorophore set based on practical performance, including brightness and stability, we developed a toolkit that supports reliable discrimination of labeled populations using microscopy or flow cytometry, even in *in vivo* competition assays. Moreover, combinatorial labeling with his set of fluorophores can resolve up to 21 distinct fluorescently labeled populations.

An important conclusion from this work and related studies is that fluorophore performance requires direct validation in the system in which probes will be used. For example, although mStayGold was expected to perform as a strong green fluorescent protein based on prior reports [24], it performed poorly in *C. albicans* under our experimental conditions. This is consistent with previous studies showing that fluorescent protein behavior is context dependent and influenced by factors such as folding, maturation, intracellular environment, expression level, and acquisition conditions [15, 30, 31]. Our results therefore underscore the importance of species-specific benchmarking when developing multiplex labeling strategies for use in a target species. In practice, the utility of a multicolor panel depends on whether individual signals remain bright, stable, and distinguishable under experimental conditions. Beyond strain discrimination, CandiChrome also proved useful for protein localization studies in *C. albicans*. Combinatorial tagging of Cox4, Efg1, Ras1 and Vph1 generated interpretable localization patterns corresponding to mitochondria, the nucleus, the plasma membrane, and vacuolar structures, respectively. This extends the value of the platform beyond competitive tracking and mixed-strain experiments and highlights its utility for combining cell biological and population-level analyses with the same fluorescent toolkit.

Several considerations in the application of CandiChrome should be noted. For example, although we identified a robust seven-fluorophore set for *C. albicans*, the resolvable number of labels are strongly influenced by instrumentation, acquisition settings, target abundance, and the biological context in which the fluorophores are used. In addition, fluorophore behavior may vary between different culture conditions, including alternative morphogenetic states, environmental stresses, and host niches. As is common in mouse experiments, mouse-to-mouse variation also contributed to variability in *in vivo* readouts, and this was evident in our pilot studies. Such variation is an inherent feature of host-associated experiments and should be taken into account in the design, replication, and interpretation of future applications of CandiChrome in animal models. More broadly, additional work will be needed to define the upper limits of multiplexing in more complex infection settings and to determine how readily this framework can be extended across strain backgrounds and other *Candida* species.

In summary, CandiChrome provides an empirically optimized multicolor labeling toolkit for *C. albicans*. By enabling strain tracking, protein localization, flow cytometric analysis, and *in vivo* discrimination of mixed populations, this platform expands the tools available for studying this species. In doing so, it also offers a practical blueprint for the development of organism-specific multiplex fluorescent systems in settings where multicolor labeling remains underdeveloped.

## Materials and Methods

### Culture conditions

*C. albicans* strains used in this study are listed in Supplementary Table S1. Unless otherwise indicated, single colonies of *C. albicans* were inoculated into 3 mL of liquid YPD medium (1% yeast extract, 2% bactopeptone, and 2% dextrose) and grown overnight at 30°C.

### Plasmid construction

To generate *pENO1*-FP-labeled *C. albicans* strains, pRB1399 was constructed to enable integration of Golden Gate Assembly (GGA) [32] constructs into the *C. albicans NEUT5L* locus. The *NEUT5L* targeting sequence, which has two *NEUT5L* segments in tandem separated by a PacI site, was PCR amplified from pRB1333 (Addgene #122378) using oligos 6060/6061. The PCR product was digested with SacI/SacII and ligated into pRB1397 (pSFS2A-GGA adapter) [33] that had also been digested with SacI/SacII to yield pRB1399.

pRB542 contains the *ENO1* promoter, a codon-optimized dTomato gene, the *TEF* terminator, and a *SAT1* selectable marker [34]. dTomato was PCR amplified from pRB542 using oligos 4568/4569 and digested with XhoI. pSFS2A [35] was linearized with XhoI, treated with calf intestinal phosphatase to dephosphorylate sticky ends, and ligated to the insert. Correct orientation of the insert was confirmed using oligos 4568/4438 to generate pRB921 (Addgene #256752).

*C. albicans* codon-optimized mStayGold and mEGFP were synthesized by BioBasic. Each fluorescent protein (FP) coding sequence was flanked by an XhoI site at the 5’ end and an FRT-BamHI sequence at the 3’ end. These constructs were cloned into pUC57 to generate pRB2288 and pRB2311, respectively. The mStayGold and mEGFP genes were then cloned into XhoI/BamHI sites in pSFS2A [35] yielding pRB2340 (Addgene #256742) and pRB2342 (Addgene **#**256743), respectively.

*C. albicans* codon-optimized mTagBFP2 was synthesized by Gene Universal in the pUC57 backbone (as pRB1203). mTagBFP2 was digested from pRB1203 using XhoI/SalI and ligated into pSFS2A[35] which had been digested using XhoI and CIP treated to generate pRB1236 (Addgene #256744). Orientation of the insert was determined by digestion with SacII/XhoI.

*C. albicans* codon-optimized mScarlet3-S2, mKate2, mAmetrine1.1, mTurquoise2, smURFP, miRFP670nano3, and BDFP1.6 were synthesized by Gene Universal. Each fluorescent protein coding sequence was flanked by an XhoI site at the 5’ end and an FRT-BamHI sequence at the 3’ end. These constructs were present in a pUC57-BsaI-free backbone, and referred to as pRB2426, pRB2427, pRB2428, pRB2429, pRB2458, pRB2459, and pRB2460, respectively. Each plasmid was digested with XhoI/BamHI and ligated into pSFS2A [35] also digested with XhoI/BamHI, yielding pRB2431 (Addgene #256745), pRB2432 (Addgene #256746), pRB2433 (Addgene #256747), pRB2434 (Addgene #256748), pRB2461 (Addgene #256749), pRB2462 (Addgene #256750), and pRB2463 (Addgene #256751), respectively.

To integrate different FPs into the *C. albicans NEUT5L* locus [29], a series of integration plasmids were generated using a GGA approach. These constructs were derived from the pRB1399 acceptor vector. Two fragments were commonly used: the *ENO1* promoter and the *TDH3* terminator were PCR amplified from genomic DNA (gDNA) of *C. albicans* SC5314 (CAY12597) using oligos 9795/9728 and 9799/9800, respectively. A third fragment was PCR amplified from pRB921 with oligos 9880/9881 for dTomato, pRB1236 with oligos 9882/9883 for mTagBFP2, pRB895 (Addgene #124586) [22] with oligos 9882/9884 for mNeonGreen, pRB2461 with oligos 9882/10055 for smURFP, pRB2432 with oligos 9882/10047 for mKate2, pRB2463 with oligos 9882/10054 for BDFP1.6, pRB2462 with oligos 9882/10049 for miRFP670nano3, pRB2340 with oligos 9882/10053 for mStayGold, pRB2433 with oligos 9882/10048 for mAmetrine1.1, pRB2434 with oligos 9882/10050 for mTurquoise2, pRB2431 with oligos 9882/10056 for mScarlet3-S2, pRB897 (Addgene #124587) [22] with oligos 9882/10051 for mScarlet-I, and pRB2342 with oligos 9882/10052 for mEGFP. The three fragments were combined by GGA with BsaI into pRB1399 to yield pRB2501, pRB2513, pRB2514, pRB2544, pRB2545, pRB2546, pRB2547, pRB2548, pRB2549, pRB2550, pRB2551, pRB2552, and pRB2553, respectively.

The *RAS1*-mNeonGreen plasmid (pRB996) was generated by PCR fusion of three fragments: the upstream *RAS1* region amplified from SC5314 genomic DNA with oligos 5088/5089, the mNeonGreen sequence amplified from pRB895 [22] with oligos 5090/5091, and the *RAS1* fragment containing downstream sequence amplified from SC5314 genomic DNA with oligos 5092/5115. The fused product was amplified with oligos 5088/5115, digested with KpnI and XhoI, and cloned into pSFS2A [35] digested with the same enzymes.

### *C. albicans* strain construction

To generate Eno1-fusion fluorescently labeled strains, FP genes were PCR amplified from the corresponding plasmids. Oligos 9547/9548 were used for all FPs except dTomato, which was amplified using oligos 9739/9548. The source plasmids were as follows: pRB1236 for mTagBFP2, pRB2434 for mTurquoise2, pRB2340 for mStayGold, pRB895 [22] for mNeonGreen, pRB2342 for mEGFP, pRB2433 for mAmetrine1.1, pRB921 for dTomato, pRB897 for mScarlet-I, pRB2431 for mScarlet3-S2, pRB2432 for mKate2, pRB2462 for miRFP670nano3, pRB2463 for BDFP1.6, and pRB2461 for smURFP. The resulting PCR products were transformed into SC5314 and selected on YPD supplemented with nourseothricin (200 μg/mL; Jena Bioscience). The corresponding Eno1-FP-expressing strains were CAY16975, CAY17099, CAY16965, CAY16971, CAY17002, CAY17060, CAY17323, CAY16999, CAY16902, CAY17058, CAY17299, CAY17295, and CAY17302, respectively. Correct integration was verified by colony PCR. The dTomato-, miRFP670nano3-, BDFP1.6-, and smURFP-labeled strains were confirmed using oligos 7764/9319, whereas all other strains were confirmed using oligos 7763/4438.

To generate p*ENO1*-FP strains at the *C. albicans NEUT5L* locus, the following plasmids were used: pRB2513 for mTagBFP2, pRB2550 for mTurquoise2, pRB2548 for mStayGold, pRB2514 for mNeonGreen, pRB2553 for mEGFP, pRB2549 for mAmetrine1.1, pRB2501 for dTomato, pRB2552 for mScarlet-I, pRB2551 for mScarlet3-S2, pRB2545 for mKate2, pRB2547 for miRFP670nano3, pRB2546 for BDFP1.6, and pRB2544 for smURFP. Each plasmid was digested with PacI and transformed into SC5314. Transformants were selected on YPD supplemented with nourseothricin. Correct integration was verified by colony PCR using oligonucleotides 4906/7832, and 4439/4907. In the same order as listed above, the corresponding strain numbers were CAY17802, CAY18103, CAY18111, CAY17804, CAY18129, CAY18106, CAY18109, CAY18127, CAY18131, CAY18114, CAY18116, CAY18126, and CAY18321, respectively.

To generate double-labeled strains, the *SAT1* marker in the initial p*ENO1*-FP transformants integrated at *NEUT5L* was recycled by growth on YP medium supplemented with 2% maltose. The resulting strains were then subjected to a second round of transformation with the p*ENO1*-FP plasmids described above, using the same transformation and selection procedure, so that a second fluorescent protein was integrated at the other *NEUT5L* allele. Thus, the final double-color strains carried one FP at each *NEUT5L* allele. The corresponding strain numbers are listed in Supplementary Table S1.

To generate a strain expressing FP fusions that localize to different cellular locations, a Cox4-dTomato fusion introduced into SC5314 by transformation with a PCR product amplified from pRB921 using oligos 10243/10244. Correct integration was verified by PCR using oligos 10245/5222. Following *SAT1* recycling, a Vph1-miRFP670nano3 fusion was introduced by transformation with a PCR product amplified from pRB2462 using oligos 10246/10247. Correct integration was verified by colony PCR using oligos 10248/4360. After an additional round of *SAT1* recycling, an *EFG1*-mTagBFP2 fusion was generated by transformation with a PCR product amplified using oligos 4446/8774. Correct integration was verified by PCR using oligos 5021/4360. Finally, the *RAS1*-mNeonGreen addback construct (pRB996) was linearized with PacI and transformed into the resulting strain. Correct integration was verified by PCR using oligos 5089/5101 to generate strain CAY19263 expressing Cox4-dTomato, Vph1-miRFP670nano3, Efg1-mTagBFP2, and Ras1-mNeonGreen.

### Fluorescence microscopy

*C. albicans* strains were grown overnight in 3 mL of synthetic complete dextrose (SCD) medium (0.7% yeast nitrogen base without amino acids, 2% glucose, 0.17% amino acid dropout mix lacking uracil, leucine, histidine, and arginine, supplemented with 0.045% uracil, 0.135% leucine, 0.045% histidine, 0.045% arginine, and 0.0025% uridine) at 30°C and visualized using an APEXVIEW APX100 microscope (Evident) or a FLUOVIEW FV5000 confocal microscope (Evident). For imaging of the multicolor strain CAY19263, cells were grown overnight at 30°C in RPMI 1640 medium supplemented with 0.165 M MOPS and 2% glucose [36] and visualized using the APEXVIEW APX100 microscope. Representative images were processed using ImageJ for uniform brightness adjustment and pseudocoloring. For strains used in fluorescence quantification, fluorescence intensity heat maps were additionally generated in ImageJ. Fluorescence intensity was quantified using CellProfiler [37] from whole-field images containing at least 100 cells per image. Cells were automatically segmented, and the average pixel intensity was measured for each cell. For photostability analysis, cells were progressively photobleached by repeated 30 ms exposures at 30% LED power using the 358-, 488-, 580-, and 650-nm excitation channels on an APEXVIEW APX100 microscope.

### Growth curve analysis

All *C. albicans* strains were grown overnight, washed twice with H_2_O, and quantified using a NanoDrop 2000c spectrophotometer (Thermo Fisher Scientific). Cells were then diluted to an OD_600_ of 0.01 in YPD medium, SCD medium, Lee’s glucose medium (17.3 g/L Lee’s powder, 1 mM MgCl_2_·6H_2_O, 0.2 µM ZnSO_4_·7H_2_O, 10 µg/mL biotin, 1.25% glucose, pH 6.8), or cecal content harvested from penicillin- and streptomycin-treated female C57BL/6J mice. Growth was monitored using an Epoch 2 microplate reader (Agilent Technologies).

### Gastrointestinal colonization model

For colonization experiments, 7-to 8-week-old female C57BL/6J mice (stock no. 000664, The Jackson Laboratory) were used in an antibiotic-treated mouse model. Mice received drinking water supplemented with penicillin (1.5 mg/mL), streptomycin (2 mg/mL), and 2.5% glucose throughout the experiment. Six fluorescently labeled *C. albicans* strains were grown overnight in 3 mL YPD, subcultured into 10 mL fresh YPD for 5 h, washed three times with H_2_O, and adjusted by OD_600_. For competition experiments, strains were mixed at equal ratios and inoculated into mice at a total inoculum of 10^8^ cells. For flow cytometric spectral unmixing, each labeled strain was also prepared and inoculated individually. Fecal pellets were collected at indicated time points, homogenized in PBS supplemented with antibiotics (500 μg/mL penicillin, 500 μg/mL ampicillin, 250 μg/mL streptomycin, 225 μg/mL kanamycin, 125 μg/mL chloramphenicol, and 125 μg/mL doxycycline), serially diluted, and plated on YPD agar.

### Flow cytometry

For flow cytometric analyses, *C. albicans* strains were grown overnight in 3 mL of SCD medium at 30°C and analyzed on a Cytek Aurora flow cytometer (Cytek Biosciences) equipped with 355-nm UV, 405-nm violet, 488-nm blue, and 640-nm red lasers. The same platform was also used for analysis of the expanded 21-sample panel (Fig. 5A), for which spectral unmixing was performed using the instrument’s built-in software to separate the different fluorescent signals.

For *in vitro* spectral unmixing analysis, *C. albicans* strains were grown overnight in 3 mL SCD at 30°C. Single-color controls and a mixed population containing all fluorescent strains were analyzed using a BD FACSymphony A5 SE flow cytometer (BD Biosciences) equipped with 349-, 404-, 488-, 561-, and 637-nm lasers. Spectral unmixing was performed using the instrument’s built-in software to verify that the fluorescent proteins could be distinguished within the mixed population.

For flow cytometric analysis of colonies recovered from fecal samples collected during *in vivo* competition experiments (Fig. 6B, C and Supplementary Fig. S4), YPD plates were incubated at 30°C for 3 days. All colonies were scraped from the agar surface, resuspended in H_2_O, diluted as needed, and analyzed using a BD FACSymphony A5 SE flow cytometer. Statistical significance was assessed by ordinary one-way ANOVA with Tukey’s multiple comparisons test.

### Automated Deconvolution and Population Quantification

To distinguish between 21 distinct cell populations (comprising 6 single-color controls and 15 dual-color combinations), we developed a computational pipeline in R (v4.3.1). The 21 reference populations served as a training set for an automated cell classification pipeline. After arcsinh transformation and downsampling, the fluorescence profiles of the reference cells were used to train a k-nearest neighbors (kNN) algorithm (k=15). This model was then applied to deconvolve the “mixed” sample. Each cell in the mixture was assigned to one of the 21 reference identities based on its proximity in the UMAP-embedded high-dimensional space. The relative proportion of each population within the mixture was subsequently calculated from the predicted classification results. The code for analysis is available on https://github.com/jcxu1999/Candida-Flow-Analysis.git.

### Ethics

Animal studies were performed according to protocols approved by the Brown University Institutional Animal Care and Use Committee (IACUC; protocol number 24-09-0007) and conducted through Brown University’s Center for Animal Resources and Education (CARE). All experiments were performed in accordance with relevant institutional guidelines and regulations, and the study was conducted in compliance with ARRIVE guidelines.

## Data availability

All data supporting the findings of this study are included in this article and its Supplementary Information files. Additional raw data are available from the corresponding author upon reasonable request.

## Acknowledgements

We would like to thank members of the Bennett lab for their advice and support throughout this project. We thank Kevin Carlson and Mark Dooner for assistance with flow cytometry analysis, and Geoff Williams and Emily Cronin-Furman for assistance with confocal microscopy. This work was supported by National Institutes of Health grants AI081704, AI141893, AI175177 and AI168222.

## Author contributions

C.-L.K. conducted most of the experiments. C.-L.K., J.X. and C.F. constructed plasmids and strains. J.X. assisted with UMAP plotting. C.-L.K. and J.X. performed confocal microscopy and *in vivo* flow cytometry. C.F. provided technical advice and guidance. R.J.B. supervised the project. C.-L.K. wrote the manuscript with input from all authors. All authors reviewed the manuscript.

