## Supplementary Information for "A CandiChrome toolkit for multicolor labeling of *Candida* cells"

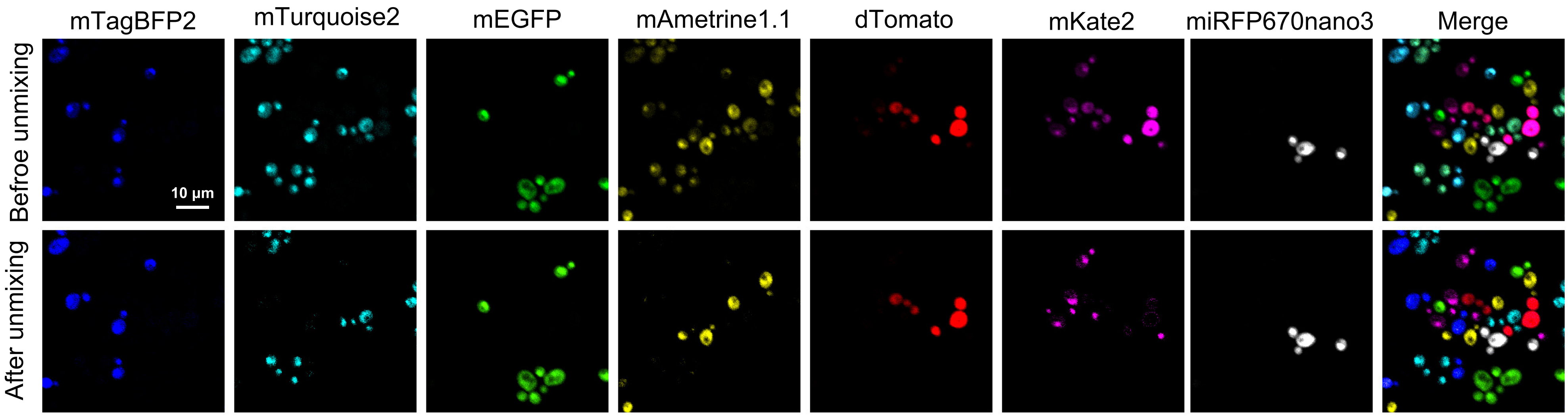


**Figure S1. Spectral unmixing of the CandiChrome panel using an Evident FV3000 confocal microscope.**
Representative fluorescence images of mixed *C. albicans* populations expressing the indicated CandiChrome fluorophores before and after spectral unmixing. Cells were grown overnight in SCD at 30 °C and imaged on an FV3000 confocal microscope. Unmixing was performed using the built-in software. Scale bar, 10 µm.

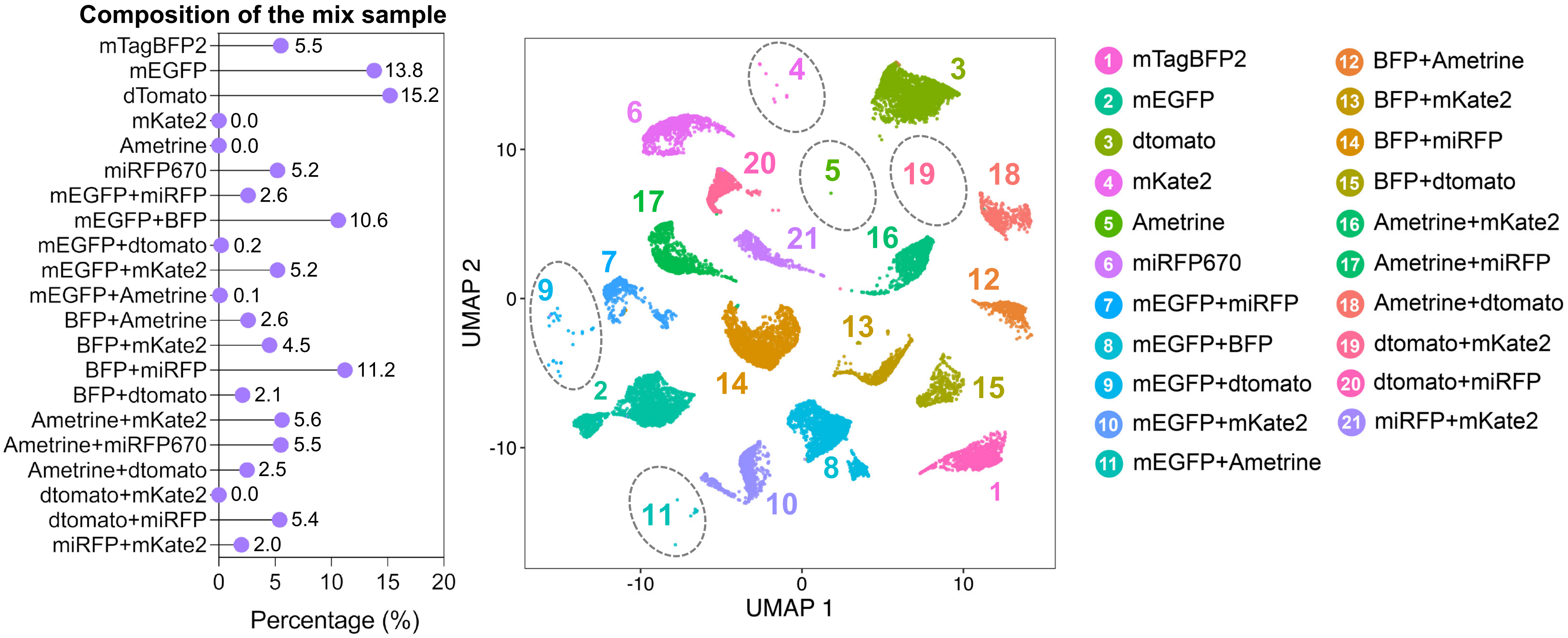


**Figure S2. Resolution of a 16-population sample by flow cytometry and UMAP.**
Composition of the mixed sample (left), UMAP projection (center), and population labels (right). Data were acquired on a Cytek Aurora flow cytometer and analyzed in R.

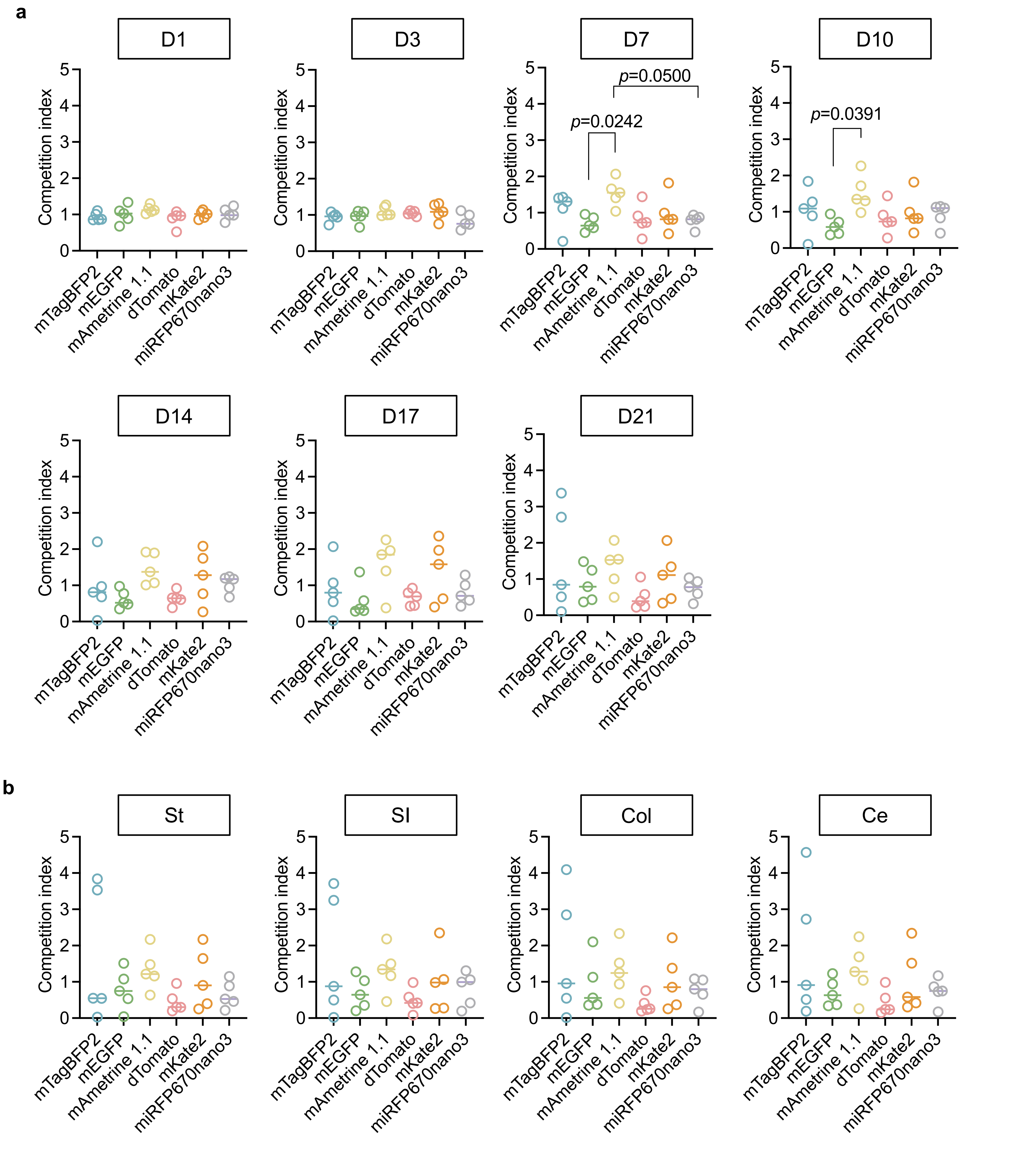


**Figure S3. Competition indices of CandiChrome-labeled strains during gastrointestinal colonization.**

**a,** Competition index values for each CandiChrome-labeled strain recovered from fecal samples at the indicated time points. Each symbol represents one mouse; horizontal lines indicate the mean. **b,** Competition indices for each strain in gastrointestinal tissues at endpoint. St, stomach; SI, small intestine; Col, colon; Ce, cecum. Each symbol represents one mouse; horizontal lines indicate the mean. Statistical significance was assessed by ordinary one-way ANOVA with Tukey's multiple-comparisons test. Exact P values are shown in the figure.

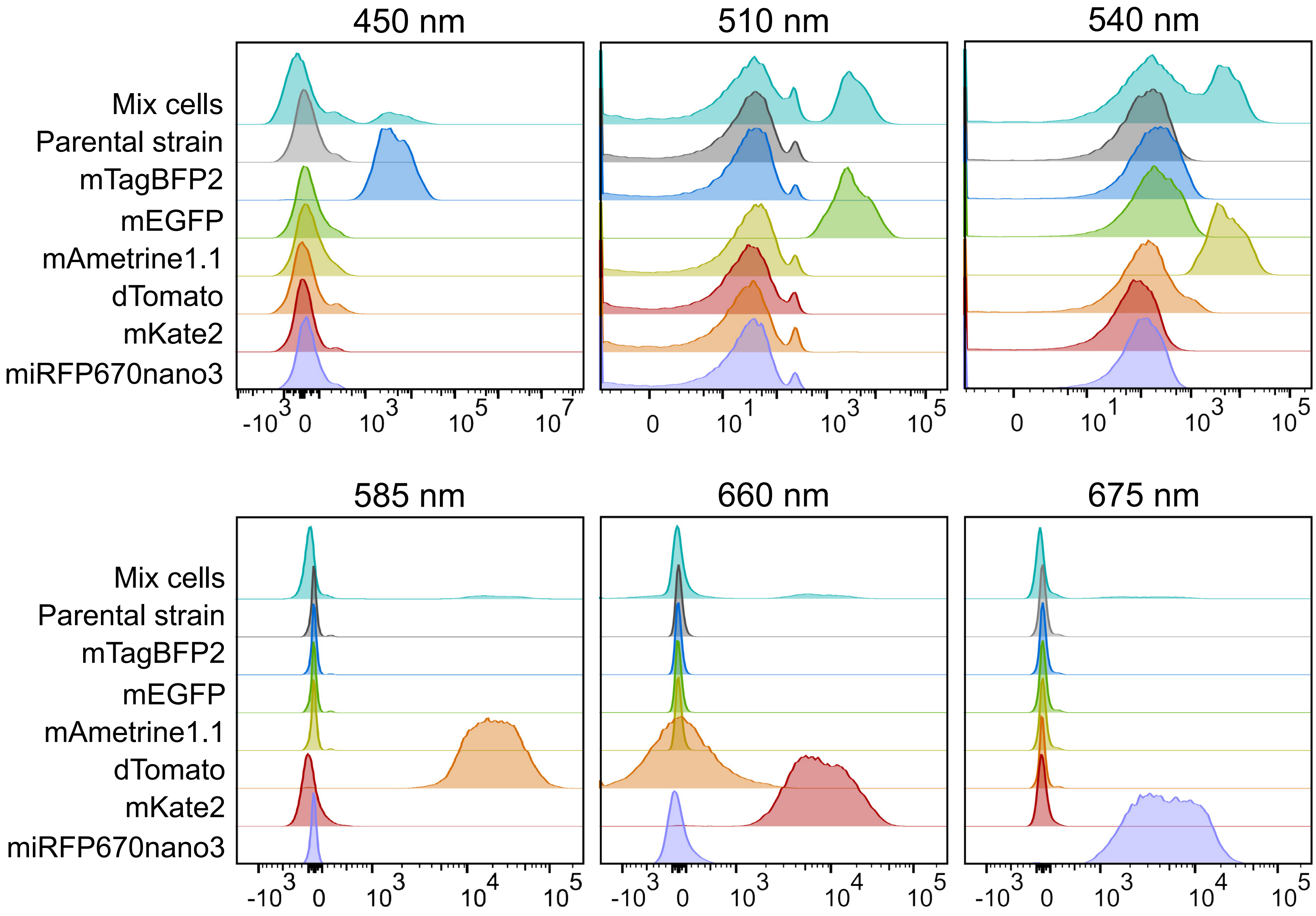


**Figure S4. Representative flow cytometry profiles of an *in vivo*-derived CandiChrome sample.**
Fluorescence intensity distributions in the indicated emission channels for a mixed-cell sample recovered from day 1, Comp5 of the *in vivo* competition assay. Individual CandiChrome strains and the parental strain (SC5314) are shown for comparison.

**Table S1. Strains used in this study.**

| Strains | Genotype | Source |
|---|---|---|
| CAY12597 | SC5314 (Wild-type) | [1] |
| CAY16975 | SC5314 Eno1-mTagBFP2 fusion | This study |
| CAY17099 | SC5314 Eno1- mTurquoise2 fusion | This study |
| CAY16965 | SC5314 Eno1- mStayGold fusion | This study |
| CAY16971 | SC5314 Eno1- mNeonGreen fusion | This study |
| CAY17002 | SC5314 Eno1- mEGFP fusion | This study |
| CAY17060 | SC5314 Eno1- mAmetrine1.1 fusion | This study |
| CAY17323 | SC5314 Eno1- dTomato fusion | This study |
| CAY16999 | SC5314 Eno1- mScarlet-I fusion | This study |
| CAY16902 | SC5314 Eno1- mScarlet3-S2 fusion | This study |
| CAY17058 | SC5314 Eno1- mKate2 fusion | This study |
| CAY17299 | SC5314 Eno1- miRFP670nano3 fusion | This study |
| CAY17295 | SC5314 Eno1- BDFP1.6 fusion | This study |
| CAY17302 | SC5314 Eno1- smURFP fusion | This study |
| CAY17802 | SC5314 pENO1- mTagBFP2 Neut5L | This study |
| CAY18103 | SC5314 pENO1- mTurquoise2 Neut5L | This study |
| CAY18111 | SC5314 pENO1- mStayGold Neut5L | This study |
| CAY17804 | SC5314 pENO1- mNeonGreen Neut5L | This study |
| CAY18129 | SC5314 pENO1- mEGFP Neut5L | This study |
| CAY18106 | SC5314 pENO1- mAmetrine1.1 Neut5L | This study |
| CAY18109 | SC5314 pENO1- dTomato Neut5L | This study |
| CAY18127 | SC5314 pENO1- mScarlet-I Neut5L | This study |
| CAY18131 | SC5314 pENO1- mScarlet3-S2 Neut5L | This study |
| CAY18114 | SC5314 pENO1- mKate2 Neut5L | This study |
| CAY18116 | SC5314 pENO1- miRFP670nano3 Neut5L | This study |
| CAY18126 | SC5314 pENO1- BDFP1.6 Neut5L | This study |
| CAY18321 | SC5314 pENO1- smURFP Neut5L | This study |
| CAY19263 | SC5314 Cox4-dTomato; Vph1-miRFP670nano3; Efg1-mTagBFP2;Ras1-mNeonGreen | This study |
| CAY18410 | SC5314 pENO1-mEGFP+ pENO1-miRFP670nano3 Neut5L | This study |
| CAY18333 | SC5314 pENO1-mEGFP+ pENO1-mTagBFP2 Neut5L | This study |
| CAY18336 | SC5314 pENO1-mEGFP+ pENO1-dTomato Neut5L | This study |

| | | |
|---|---|---|
| CAY18335 | SC5314 pENO1-mEGFP+ pENO1-mKate2 Neut5L | This study |
| CAY18334 | SC5314 pENO1-mEGFP+ pENO1-mAmetrine1.1 Neut5L | This study |
| CAY18407 | SC5314 pENO1-mTagBFP2+ pENO1-mAmetrine1.1 Neut5L | This study |
| CAY18342 | SC5314 pENO1-mTagBFP2+ pENO1-mKate2 Neut5L | This study |
| CAY18344 | SC5314 pENO1-mTagBFP2+ pENO1-miRFP670nano3 Neut5L | This study |
| CAY18405 | SC5314 pENO1-mTagBFP2+ pENO1-dTomato Neut5L | This study |
| CAY18352 | SC5314 pENO1-mAmetrine1.1 + pENO1-mKate2 Neut5L | This study |
| CAY18350 | SC5314 pENO1-mAmetrine1.1 + pENO1-miRFP670nano3 Neut5L | This study |
| CAY18341 | SC5314 pENO1-mAmetrine1.1 + pENO1-dTomato Neut5L | This study |
| CAY18404 | SC5314 pENO1-dTomato + pENO1-mKate2 Neut5L | This study |
| CAY18409 | SC5314 pENO1-dTomato + pENO1-miRFP670nano3 Neut5L | This study |
| CAY18338 | SC5314 pENO1-miRFP670nano3 + pENO1-mKate2 Neut5L | This study |

1. **Bennett RJ, Johnson AD.** 2006. The role of nutrient regulation and the Gpa2 protein in the mating pheromone response of *C. albicans*. Mol Microbiol **62:** 100-119.

**Table S2. Oligonucleotides used in this study.**

| Number | Sequence (5’->3’) |
|---|---|
| 6060 | GGACCGCCGCGGTAAACAAGTGGTATTCAAGCACAAT |
| 6061 | GGACCGGAGCTCCAGGAAGGACGATGAAGGA |
| 4568 | GGACCGCTCGAGCGGATCCCCGGGTTAATT<br>AACGGTATGGTTTCAAAAGGTGAAGAAGTT<br>ATTAAAG |
| 4569 | GGACCGCTCGAGTTATTTATACAATTCATC<br>CATACCATACAAAAACAA |
| 4438 | CTCAACCATAGCAATCATGG |
| 9795 | GGACCGGGTCTCTCAGTCATTTGTATCTTTAGTAGACATGATTGT |
| 9728 | GGACCGGGTCTCTCATTGTTGTAATATTCCTGAATTATCAAT |
| 9799 | GGACCGGGTCTCTTAAAGTAAAACCAGACTTTGATTTGATT |
| 9800 | GGACCGGGTCTCTAGTATGGTAATAGGAAGTCAAAAGAAAGA |
| 9880 | GGACCGGGTCTCTAATGGTTTCAAAAGGTGAAGAAGTTATTAAAG |
| 9881 | GGACCGGGTCTCTTTTATTTATACAATTCATCCATACCATACAAAAAC |
| 9882 | GGACCGGGTCTCTAATGGGTGGTAGTGGTATGGTTTC |
| 9883 | GGACCGGGTCTCTTTTAGTTCAATTTATGACCTAATTTTGATGG |
| 9884 | GGACCGGGTCTCTTTTATTTATACAATTCATCCATACCCATAACA |
| 10055 | GGACCGGGTCTCTTTTATGACATAGCTTTAATAATATAATCAAAATATGGA<br>G |
| 10047 | GGACCGGGTCTCTTTTACTATCTATGACCTAATTTTGATGGTAAATCA |
| 10054 | GGACCGGGTCTCTTTTAAATTTCAGTTTCTGAAATAACTCTAGC |
| 10049 | GGACCGGGTCTCTTTTATGATTGTTGAATAGCAATACCCA |
| 10053 | GGACCGGGTCTCTTTTATTATAAATGAGCTTCTAAAGTTTCAGATTG |
| 10048 | GGACCGGGTCTCTTTTACTATTTATATAATTCATCCATACCTGGAGTA |
| 10050 | GGACCGGGTCTCTTTTACTATTTATATAATTCATCCATACCTAAAGTAATA<br>CCA |
| 10056 | GGACCGGGTCTCTTTTACTATGAACCACCTGAACCAC |
| 10051 | GGACCGGGTCTCTTTTACTAATATAATTCATCCATACCACCTG |
| 10052 | GGACCGGGTCTCTTTTATTATTTATATAATTCATCCATACCTAAAGTAATA<br>CCA |
| 9547 | ATCTTGAGAATCGAAGAAGAATTAGGTTCTGAAGCTATCTACGCTGGTAA<br>AGATTTCCAAAAGGCTTCTCAATTGGGTGGTAGTGGTATGGTTTCTAAAG |
| 9548 | TTTGACTGCAGCTCAGTGATTAAGAGTAAAGATGGGTAAAAAATTATCAT<br>TTAATTAGTTCATATATTCAAGATGGCGGCCGCTCTAGAACTAGTGGATC |

| | |
|---|---|
| 9739 | AGAATCGAAGAAGAATTAGGTTCTGAAGCTATCTACGCTGGTAAAGATTT CCAAAAGGCTTCTCAATTGGGTGGTAGTGGTATGGTTTCAAAAGGTGAAG |
| 7764 | GTGATGATTTGACTGTCACTAACC |
| 9319 | GTGACTCCATCACCCAGTTT |
| 7763 | GATGCTTGGGTCCACTTCT |
| 4906 | ACTTTAATCCTTAGTCTTCTATTCTGAGAT |
| 7832 | ACGCAATTAATGTGAGTTAGC |
| 4439 | GCGAAAAAGTGGGCACTAAG |
| 4907 | CAAATTCATTGGAGCGATATCG |
| 10243 | AAAGTCTCTAGATGCTGGCAATGTGGTACTGTCTTGAAGGCCAAATACTT GGGTGAACCAGGAATGGCTCATCATGGTGGTAGTGGTATGGTTTCAAAAG |
| 10244 | TAGTAGAGTTCGCCAAAAGAGAACCATGAAATATATATAAAAAATAGAA TATAGCAATTGTTGGGGGTCTGTCGGGCGGCCGCTCTAGAACTAGTGGAT C |
| 10245 | CTTTGATTGGTCCTGGTGCT |
| 5222 | CTTCACCTTCAATTTCAAATTCATGAC |
| 10246 | ATGTCGAAATATTTTGAAGGTGGTGGTTCTGCTTTTGAACCATTTACTTTT AAAGGTTTATTAGACAGTGTTTTAGGTGGTAGTGGTATGGTTTCTAAAG |
| 10247 | ATGTTTAATGTTTAATGTTTATTATATATATTCAAATAAATATAATTCTTAA TTCTTTTTTATAAATTAATTGAGGCGGCCGCTCTAGAACTAGTGGATC |
| 10248 | TAATGCATTTGGTCCCACCG |
| 4360 | GTGGTTTCAGTGGCTACAAC |
| 4446 | CAAGGTTCAGTTCACCCTTCACCCCAACAACATCAAGCTAATCAATCAGC TAGCACTGTTGCCAAAGAAGAAAAGGGTGGTAGTGGTATGGTTTCTAAAG |
| 8774 | CTTTCCAATCATTTGTTAATGAAATATATGCTATAATCTAATTTGGAATTT ATGGCAGAAAGCAGAAGGTGATGTACACCGGCCGCTCTAGAACTAGT |
| 5021 | AGCGGTAATGGGAACAGTATA |
| 5088 | GGACCGGGTACCGCATAATCTCTTTAAACT |
| 5089 | ATTATCTTCTTCACCTTTAGAAACCATGGT |
| 5090 | ATGGTTTCTAAAGGTGAAGAAGATAAT |
| 5091 | TTTATACAATTCATCCATACCCATAACATCA |
| 5092 | TGATGTTATGGGTATGGATGAATTGTATAA |
| 5115 | GGACCGCTCGAGGTTAGTTTAAGATTACAA |
| 5101 | TTGTCTTGGGAATGTAAATTTAAATGAAT |
